# Can rat cortical cultures predict what comes next, unpredictable or learned?

**DOI:** 10.64898/2026.11.16.767134

**Authors:** Cedric Cannard, William Reid

**Affiliations:** Swartz Center for Computational Neuroscience, University of California San Diego, 9500 Gilman Drive, La Jolla, CA 92093, USA; Valis Corporation, Encinitas, CA, USA

**Author notes:** Corresponding author; Lead contact: Cedric Cannard.

**Keywords:** predictive processing, anticipatory activity, rat cortical cultures, high-density multi-electrode array, stimulation artifacts, held-out validation

## Abstract

In humans, physiological activity recorded before an unpredictable stimulus reportedly varies slightly with what the stimulus will be, an effect termed predictive anticipatory activity. Whether reduced neural preparations show it is unknown. We re-analyzed a public dataset in which twelve dissociated rat cortical cultures on high-density multi-electrode arrays performed a hidden-source inference task (DANDI 001611: 27 experiments, 691,200 trials). Two features of the dataset limit what it can test. Each experiment replays one fixed 256-trial source sequence in all 100 sessions, so the upcoming source is unpredictable only in session 1. One stimulation volley per trial arrives on an exact 1-s clock, so every pre-stimulus window lies in the previous trial’s after-effects. Two questions were fixed before a confirmatory run on held-out experiments. Q1 asked whether, in session 1, a stimulation-free 250-ms window before each trial’s volley carries information about the upcoming sources beyond what trial history explains. Q2 asked whether this information grows over the 100 repetitions, as learned expectation would predict. Q1 (13 experiments, 9 cultures): decoding above a shifted-label null, ΔAUC +0.003, 95% CI -0.026 to +0.033, randomization p = 0.73, equivalent to zero within ±0.059 (the human effect expressed as AUC) and narrowly within ±0.03. Q2 (12 experiments, 8 cultures): growth ΔG +0.004, 95% CI -0.006 to +0.013, p = 0.29, equivalent within ±0.02. Pre-stimulus activity thus carried no detectable information about the stimulus, and none emerged with repetition.

**Highlights:**

- Public rat cortical culture data were re-analyzed under a plan fixed in advance
- Pre-stimulus activity held no detectable information about the coming stimulus
- This information did not grow over 100 repetitions of the same sequence
- A 1-s stimulation clock explains the apparent pre-stimulus structure in the data

**eTOC blurb:** Cannard and Reid test whether rat cortical cultures show activity that predicts an upcoming, unpredictable stimulus. Under a plan fixed in advance, pre-stimulus activity carries no detectable information about the stimulus and gains none over 100 repetitions. The 1-s stimulation clock explains the apparent pre-stimulus effects.

## Introduction

The brain is often described as a prediction machine: sensory systems learn the statistics of their inputs, form expectations, and update them when the expectations are violated^1,2^. A smaller and contested literature goes further. It reports that physiological measures in humans, such as skin conductance, heart rate and the electroencephalogram (EEG), differ in the seconds before a stimulus according to what the stimulus will be, even when its identity or timing cannot be predicted. This effect has been termed predictive anticipatory activity^3^. The pooled effect is small (standardized mean difference about 0.21), and whether it reflects methodological artifacts or genuine anticipation is still debated^3,4,5^. One way to test it is to ask whether the effect appears in reduced preparations. If anticipation of this kind depends on the recurrent, whole-brain organization of an intact and embodied cortex, it should be weaker or absent in isolated neural tissue.

Dissociated cortical cultures and human brain organoids^6,7^ are such preparations: living neural networks grown in a dish, recorded and stimulated through multi-electrode arrays (MEAs), grids of electrodes on which the tissue grows. Several stimulation experiments on them are now public. The largest we found is a hidden-source inference task in rat cortical cultures^8^. On each of 256 trials, two binary sources are drawn at random, and the culture receives a noisy report of them as one stimulation volley (none when both sources are 0); trials follow each other on a fixed 1-s grid. Two features of this dataset decide what it can test. Each experiment replays the same 256-trial sequence in all of its sessions, so a draw is unpredictable only the first time the culture meets it. And with one volley per second, every window before a stimulus falls within the after-effects of the previous one. A second public stimulation dataset, a dose-response series in human organoids recorded with shell-shaped electrode arrays^9^, has a single stimulus type delivered at scheduled times, so it cannot test whether activity predicts what a stimulus will be. That organoids respond to electrical stimulation in a parameter-dependent way is established by independent work^10,11^.

Here we re-analyze all deposited data from the hidden-source experiment and ask two questions. Does activity before an unpredictable stimulus predict what that stimulus will be? And does that information grow as the same sequence repeats? Both questions and their full analysis plan were specified and frozen (local git tag prereg-v1, 2026-10-28 16:19 PDT) before the confirmatory run on the held-out experiments, and registered as a time-stamped pre-analysis plan on the Open Science Framework during the held-out run (19:51 PDT, before any per-experiment output was written; embargoed during peer review and made public at acceptance; STAR Methods, Freeze and run timeline). The data were public and had already been analyzed, by the original authors and in our own earlier work, so this is a pre-analysis plan rather than a pre-data registration: the plan was frozen against a held-out split of the data, not against unseen data. We find no detectable predictive anticipatory activity for either question. We then show that the timing of the paradigm explains every apparent pre-stimulus effect we examined in this deposit, and we compare task activity with the stimulation-free activity recorded before each session’s first trial. Because equivalence tests bound the null, the result also defines what a decisive in-vitro experiment would need.

## Results

### The task, and two features of the deposit

In the hidden-source experiment^8^, twelve dissociated rat cortical cultures were recorded on high-density MEAs in 27 experiments of 100 sessions, each of 256 trials (691,200 trials in total). On every trial two hidden binary sources, s¹ and s², are drawn, and the deposit records both of them and the stimulation pattern delivered. We verified two features of the files on all 2,700 sessions; together they set what the data can test (STAR Methods).

First, each experiment replays one fixed 256-trial sequence of the two sources in all 100 sessions, and no two experiments share a sequence. Within a sequence the draws behave like independent fair coin flips: a pre-specified randomness audit at the level of trial positions found no departure in any sequence (smallest Holm-corrected p over 432 tests, 0.80). One pooled exploratory statistic does depart, a weak lag-8 autocorrelation of s¹ (Methods S1.4). The upcoming draw is therefore unpredictable to the culture only on first exposure, in session 1. From session 2 on, the draw is fixed by trial position, and the culture has already received it at that position. A test of anticipating an unpredictable draw must therefore use session 1. Later sessions test learned expectation, which is a different and conventional capacity.

Second, each stimulated trial receives a single stimulation volley of about 12 ms, on an exact 1-s clock, and nothing else stimulates the culture between volleys (Figure 1). Trials in which both sources are 0 (the (0,0) category, called no-observation trials here) receive no volley. The response evoked by a volley decays over several hundred milliseconds (Figure 1B), but the previous trial still shapes the next window: the window rate after a volley, as a ratio to the rate after a no-observation trial, ranges from 0.07 to 3.4 across held-out experiments, so some cultures are suppressed after a volley and others keep firing. Every pre-stimulus window in these data therefore lies within the previous trial’s after-effects, and a window placed more than 1 s before a trial lies before an earlier trial’s volley. This matters for our own earlier, exploratory analysis of the deposit (Methods S1.1). Its 3.45-3.10 s window ends 95 ms before the volley of trial n-3, so it tested whether activity before trial n-3 predicts the category of trial n. The bursts of synchronized activity it found at fixed offsets (0.90, 0.50 and 2.45 s before each trial’s recorded start) are responses to earlier volleys on the same grid: 0.90 s before a trial is about 100 ms after the previous volley, on the evoked network burst, and 0.50 and 2.45 s fall in the decaying tails of earlier responses (Methods S1.1). They are not stimulation events, and they also appear in no-observation trials because the neighboring trials’ volleys fall on the same grid. Most detections during the task fall inside volley spans: 89 percent in the audited experiment (80-96 percent per session; Methods S1.2), with a share that varies between experiments. Rates that include these artifacts (about 4.8-6.4 detections per second per channel) therefore overstate spontaneous activity, which in the audited experiment is about 0.7 per second per channel outside artifacts and 0.03 before the first trial.

**Figure 1.**
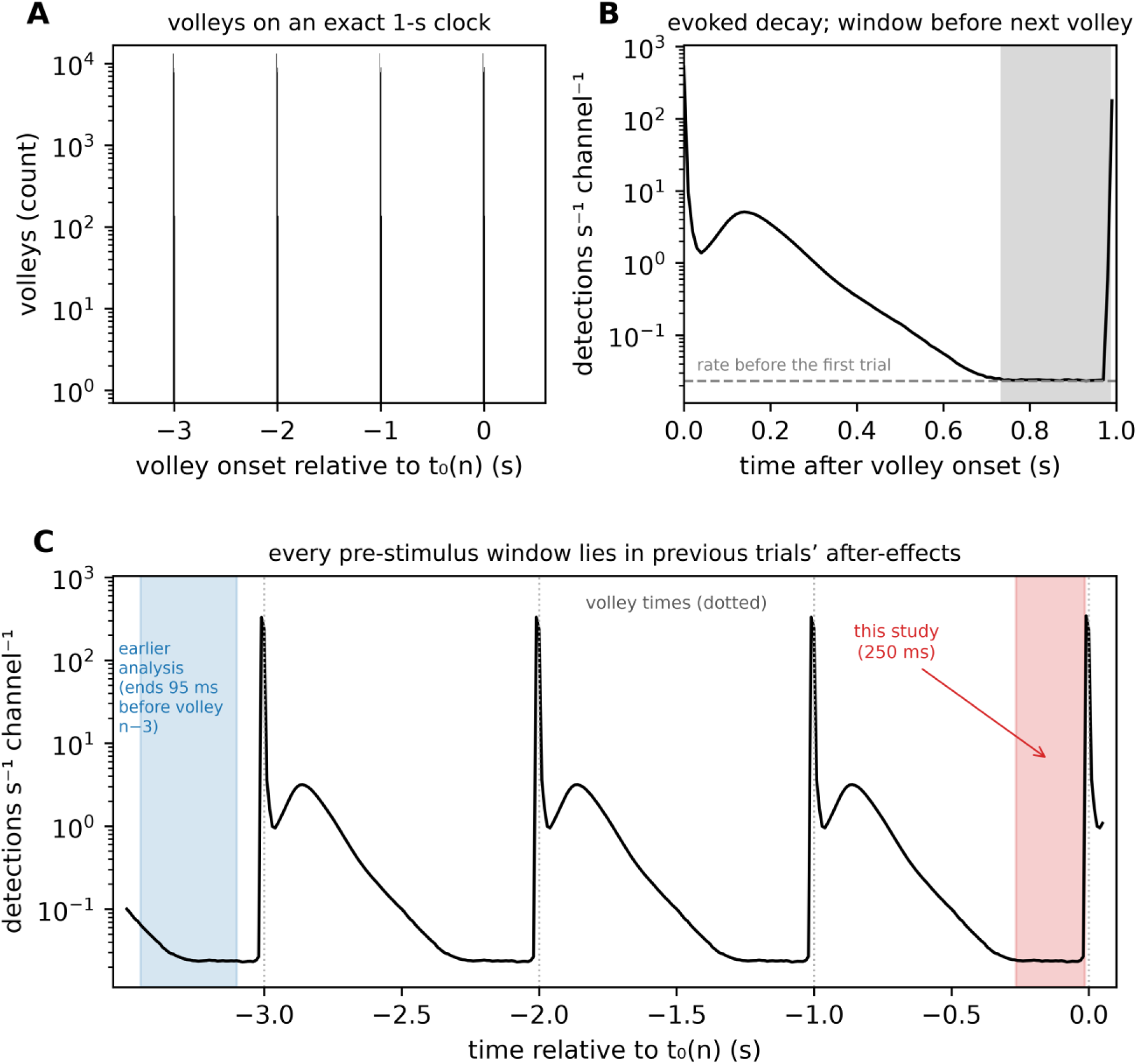
One stimulation volley per trial on a 1-s clock, and the pre-stimulus window. All panels show one experiment (C03_E02, 100 sessions). (A) Detected stimulation volleys relative to each trial’s anchor t₀(n), the fitted 1-s grid (STAR Methods), over all trials (log scale): volleys fall at exactly 0, -1, -2 and -3 s and nowhere between. (B) Detection rate per channel after a volley (log scale), with the rate before the first trial of each session (dashed). The evoked response, including a network burst that peaks about 140 ms after the volley, decays over several hundred milliseconds; the next volley arrives at 1 s. The shaded band is the 250-ms analysis window, [t₀ - 265, t₀ - 15] ms, which is stimulation-free. In this experiment the window rate is close to the rate before the first trial and far below the evoked response; across held-out experiments the previous volley still changes the window rate (Results), and this carryover is removed statistically. (C) Detection rate over the 3.5 s before t₀, with the window of this study (red) and the 3.45-3.10 s confirmatory window of our earlier analysis (blue), which ends 95 ms before the volley of trial n-3. The fixed offsets of that analysis, 0.90, 0.50 and 2.45 s before t₀, fall on responses to earlier volleys on the same grid. See also Table S1.

### No information about the upcoming draw precedes the stimulus (Q1)

Before any outcome analysis, the experiments were split into a development set, used to build and test the pipeline, and a held-out set, analyzed once in the confirmatory run (STAR Methods). For Q1 we asked whether spike counts in a stimulation-free 250-ms window ending 15 ms before each trial’s own volley predict the trial’s two hidden sources, in session 1 of each held-out experiment (13 experiments in 9 cultures). Because the window carries the after-effects of earlier trials, we first removed what trial history explains (the previous labels, the previous stimulation and the previous evoked response), fitting this correction on training data only. A classifier then decoded each source from the remaining activity, scored by the area under the receiver operating characteristic curve (AUC; 0.5 is chance, 1 is perfect). Rather than assume that chance is exactly 0.5, we compared each experiment’s AUC with a null distribution built by circularly shifting its label sequence, which keeps the structure of the activity and of the labels but breaks their alignment (the sequence-shift null). ΔAUC is the observed AUC minus the mean of that null. Experiments were pooled with a random-effects model, experiments nested in cultures, fit by restricted maximum likelihood (REML); the primary p-value comes from a randomization test that refits this model to draws from each experiment’s null. Pooled decoding was at chance: ΔAUC +0.003 (SE 0.013; 95% CI -0.026 to +0.033; randomization p = 0.73; REML p = 0.80) (Figure 2).

**Figure 2.**
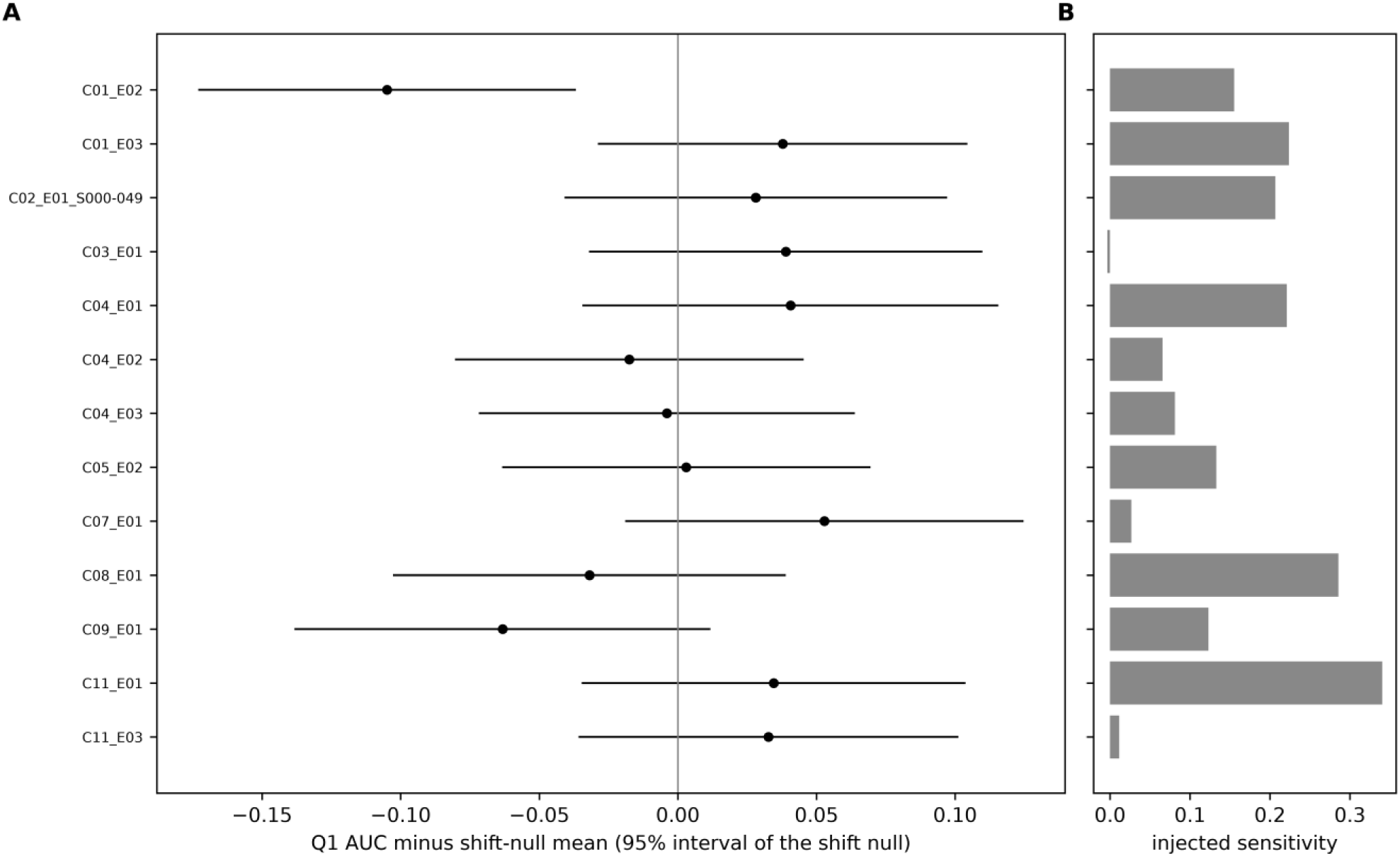
No pre-stimulus information about the upcoming draw in session 1 (Q1). (A) For each of the 13 held-out experiments (9 cultures), the session-1 decoding AUC of the two hidden sources from history-residualized window features, minus the mean of the experiment’s sequence-shift null; bars are the 95% interval of that null (225 shifts). Pooled estimate +0.003 (95% CI -0.026 to +0.033; randomization p = 0.73), equivalent to zero within ±0.059 and, narrowly, within the secondary ±0.03. (B) Each experiment’s injected sensitivity, the ΔAUC produced by a synthetic 5% multiplicative rate change on all included channels with permuted labels; three experiments (C03_E01, C11_E03, C07_E01) are nearly blind at that size. C02_E01_S000-049 is the held-out half (sessions 1-50) of a culture recorded in a single experiment. See also Tables S2 and S3.

A non-significant result is not evidence of absence, so we also tested equivalence^12^: two one-sided tests (TOST) against the smallest effect of interest fixed before the run, ±0.059, which is the pooled human effect^3^ expressed as AUC. The estimate is equivalent to zero within this bound (TOST p = 0.001; 90% CI -0.021 to +0.027) and, narrowly, within the secondary bound of ±0.03 (TOST p = 0.037; the 90% upper limit of 0.0275 lies just inside). A Bayes factor, the evidence for an effect relative to the evidence for none (BF10; values below 1 favor the null), agrees: BF10 = 0.17 with a Cauchy prior scaled to the smallest effect of interest, and 0.31 and 0.09 at half and twice that scale. Under the decision rule fixed before the run (STAR Methods), this is outcome A, a bounded null at ±0.059. Between-culture variance was estimated at zero. Between-experiment variance (τ² = 0.0009) comes mostly from one experiment, C01_E02, recorded at 13 days in vitro (DIV 13), whose AUC falls below its null (ΔAUC - 0.105); without it, the pooled estimate is about +0.012 and the outcome is unchanged (Methods S1.9). A pre-specified exploratory check that decoded the (0,1) versus (0,0) categories in session 1, and could not change either outcome, was likewise null (+0.013, 95% CI -0.032 to +0.059, p = 0.52; Methods S1.10). Two post hoc descriptions lie outside the registered decision rule. The pooled effect corresponds to a standardized effect dz = 0.01 (Hedges g = 0.01 after small-sample correction at df = 8; ΔAUC = Φ(dz/√2) - 0.5). A flat-prior 95% interval (estimate ± 1.96 SE), -0.022 to +0.029, is reported alongside the REML t-interval; it is slightly narrower because it uses the normal rather than the t quantile.

Pre-specified sensitivity analyses, reported without testing, agree: experiments at DIV ≥ 15 only (8 experiments, 6 cultures) +0.014 (95% CI -0.017 to +0.046); a 360-ms window +0.009 (-0.014 to +0.032; randomization p = 0.35); features without history removal +0.007 (-0.016 to +0.031); rank-transformed features +0.013 (-0.015 to +0.041); all channels +0.002 (-0.027 to +0.032). A second null, which pairs each experiment’s windows with another experiment’s label sequence (the sequence-swap null), is centered on the shift null (swap minus shift mean -0.002, SD 0.013 over experiments).

### Decodability does not grow over 100 repetitions of the sequence (Q2)

If the cultures learned the repeated sequence, decoding of the upcoming label after history removal should improve across sessions. We grouped the 100 sessions of each experiment into ten blocks of ten and measured growth as G, the fitted change in block AUC from the first to the last block, again relative to a sequence-shift null (ΔG). Over the 12 held-out experiments in 8 cultures, the pooled growth was ΔG +0.004 (SE 0.004; 95% CI -0.006 to +0.013; randomization p = 0.29; REML p = 0.42) (Figure 3), equivalent to zero within the ±0.02 bound fixed before the run (TOST p = 0.003; 90% CI - 0.004 to +0.012; BF10 = 0.23). The secondary contrast, block 10 minus block 1, was +0.006 (-0.007 to +0.018). This is again outcome A, a bounded null at ±0.02. At DIV ≥ 15 (7 experiments, 5 cultures) the growth was +0.005 (-0.013 to +0.023), and with the 360-ms window +0.001 (-0.007 to +0.009). Post hoc, outside the decision rule: dz = 0.01 (Hedges g = 0.01 at df = 7, from the conversion used for Q1) and a flat-prior 95% interval of -0.005 to +0.012.

**Figure 3.**
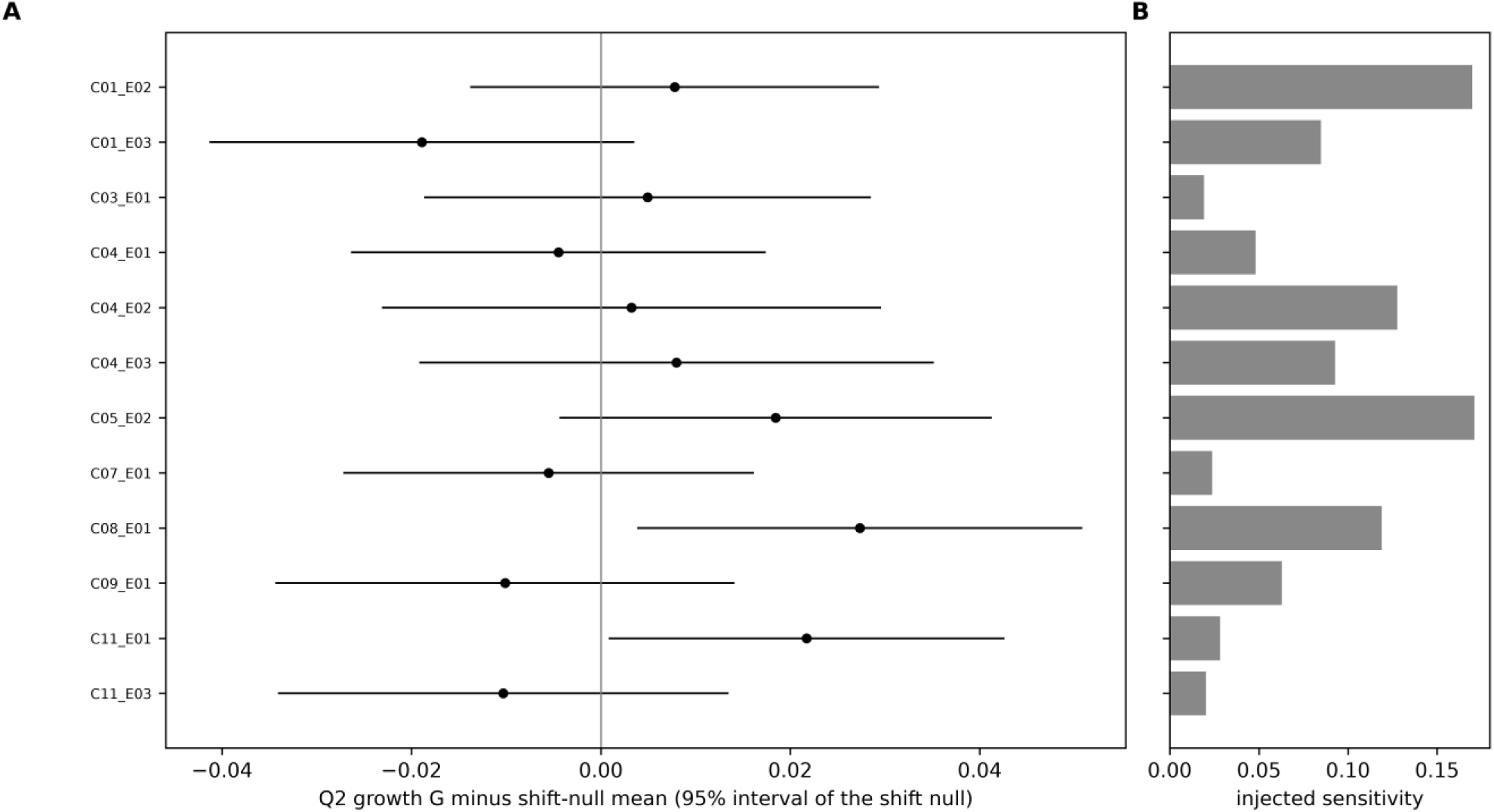
Residualized decodability does not grow over 100 repetitions of the sequence (Q2). (A) For each of the 12 held-out experiments (8 cultures), the growth statistic G (fitted change in block AUC from sessions 1-10 to sessions 91-100, history-residualized, per-channel features) minus the mean of its shift null, with the null’s 95% interval. Pooled +0.004 (95% CI -0.006 to +0.013; randomization p = 0.29), equivalent to zero within ±0.02. (B) Injected sensitivity, the ΔG produced by a synthetic rate change ramping from 0 to 10% over the 100 sessions. See also Tables S2 and S3.

The windows do change across sessions: the imprint of the previous trial’s response grows. Without history removal, the AUC for the previous label s(n-1), averaged over the 12 experiments, rises from 0.574 in block 1 to 0.612 by block 6 and stays near 0.60 afterwards (per-experiment growth +0.028, SD 0.052). After history removal, windows on the development split could not decode s(n-1); a check that had to pass before the held-out run (pooled excess +0.0005, 95% CI -0.004 to +0.005). The Q2 null therefore concerns information about the upcoming label, not the growing imprint of the previous one. A second pre-specified exploratory check asked whether the pre-stimulus category contrast mirrors the post-stimulus one across experiments (Spearman correlation). It was null (rho = 0.09, sign agreement 8 of 13, shift-null p = 0.77; Methods S1.10).

Two pre-specified interpretation controls test growth in decoding of labels that no biological process should predict, those of the trials 5 and 10 positions ahead (k = 5 and k = 10). The k = 10 control was null (+0.0005, -0.008 to +0.009). The k = 5 control was flagged: +0.0105 (95% CI +0.002 to +0.019), randomization p = 0.0025, positive in 10 of 12 experiments. Under a plan fixed before it ran, it did not replicate on the independent development split (11 experiments, 8 cultures: +0.0035, 95% CI -0.006 to +0.013, p = 0.33), and the 27 stimulation sequences share no lag-5 structure. The pre-stated rule therefore classifies it as a chance finding, but it remains unexplained. It stays nominally significant after Holm correction over 14 tests (p = 0.035; the 2 pre-specified growth controls plus 12 lag tests, a family defined after the flag) and over the two growth controls alone (Holm p = 0.005). The development interval also contains the held-out value, so the failure to replicate is weak evidence. Leave-one-out shows the effect spread across experiments; only dropping culture C04 widens the CI to include zero. It does not change the Q2 bound (Methods S1.3). The weak lag-8 autocorrelation of s¹ noted above bears only on this control (Methods S1.4).

### Controls and what the bound means

The alignment and lag-profile controls (Table S5) passed; the timing-anchor control was flagged under its literal rule. Decoding a trial’s category from counts 25-300 ms after its volley peaks at the trial itself (lag 0) in all 32 held-out tests (AUC 0.61-0.98), so labels and timestamps are aligned. Decoding the label of trial n+k from the window of trial n, on real development labels (Table S1), shows the expected carryover from previous trials (ΔAUC +0.044 at k = -1 for Q1 and +0.078 for the Q2 block AUC, without history removal) and no decoding of labels 5 to 10 trials ahead (smallest Holm p 0.98). Anchoring the windows on the recorded trial start times instead of the fitted 1-s grid changed a few window counts (32 on development, 26 on held-out data) and moved no experiment’s AUC by more than 0.0027 on development and 0.00065 on held-out data (0.08 and 0.017 of the per-experiment null SD). The pre-specified rule flags any difference; this one is the known one-sample, label-dependent jitter of the recorded start times, which the grid anchor exists to remove. Naive cross-validation by session decodes permuted labels of a fixed sequence, because the decoder can learn trial position instead of the label; position-aligned cross-validation does not. Preprocessing removed no analyzed trial.

Neither planned positive control was usable (Discussion), so sensitivity was calibrated by synthetic injection only. We added synthetic spikes that raise a channel’s window count by a fraction δ for each active source, on development data with permuted labels, and counted how often the pipeline detected them. An effect of the same size in every experiment is detected 80% of the time at ΔAUC 0.031 for Q1 (14 development units; an analysis unit is an experiment, or one half of a culture recorded in a single experiment) or 0.036 (a design like the held-out one, 9 cultures), and at a growth of 0.013 for Q2, with false-positive rates of 0.00-0.03 at nominal 0.05 (Table S2). When the same rate change is injected into all channels, its effect on decoding differs between experiments, and 80% detection needs δ ≈ 0.044-0.050 (pooled ΔAUC 0.054-0.063) for Q1 and a ramp to δ ≈ 0.09 (ΔG 0.050) for Q2; at the Q2 smallest effect of interest, power against such rate-proportional growth is only about 0.2. In the held-out experiments themselves, sensitivity to injection varies (Figures 2B and 3B; Table S3). At a 5% rate change, 10 of 13 Q1 experiments respond with ΔAUC +0.07 to +0.34, but three (C03_E01, C11_E03, C07_E01) respond with only -0.004 to +0.027 and are nearly blind at that size; the same three are the least sensitive to a 10% ramp in Q2. Each carries about the same weight in the pooled estimate as the others.

The result is therefore a bounded null for both questions at the stated sensitivity. In these cultures, activity in the 250 ms before an unpredictable stimulus carries no detectable information about that stimulus beyond what earlier trials explain. Nor does this pre-stimulus label information, our pre-specified measure of learned expectation, grow detectably over 100 sessions, even though the imprint of the previous response does. The pipeline would have detected a count change of a few percent in most experiments. This sensitivity rests on injected signal, not on a demonstrated capacity of the cultures (Discussion).

### The 1-s rhythm is task-locked, and evoked responses depress within sessions

Two descriptive analyses, outside the registered plan, use the rat recordings themselves as controls. First, we measured the normalized autocorrelation of population spiking before the first trial and in an equally long segment beginning at the first trial. The pre-task segments are short (median 3.18 s, range 2.71-4.08 s), but all 2,700 sessions contain at least 20 spikes (median 234; minimum 23). Mean autocorrelation at a 1-s lag was -0.021 before task onset and 0.357 in the matched task segments (12 cultures). The task-segment value is the stimulation clock itself, because volley detections were kept in that segment; with the 10-ms bins within -16 to +12 ms of each trial time masked, the culture mean fell to -0.03 (range -0.15 to 0.14). Before task onset the 1-s autocorrelation was at or below zero in every culture (-0.09 to 0.00). Segments this short can therefore resolve the stimulation clock, although they cannot resolve the evoked response alone. Together with the no-observation trials, this ties the 1-s structure to the task rather than to a spontaneous culture rhythm.

Second, we asked whether the evoked response clears before the next volley and whether it changes with repeated stimulation. At 600-900 ms after a volley, the difference in firing rate between stimulated and no-observation trials had a median of 0.02 spikes/s/channel across cultures (range - 0.25 to 0.64), compared with a median evoked rate of 5.83 spikes/s/channel at 100-200 ms. The evoked transient is therefore largely over before the next volley, although residual late activity varies between cultures. Within sessions, the evoked response in the second half of the 256 trials was lower than in the first half in 25 of 27 experiments (median second-half/first-half ratio 0.54, range 0.15-1.07). This short-term response depression is consistent with repeated-stimulation effects in dissociated cortical cultures^13,14^. It concerns response magnitude, not the category information tested by Q1 and Q2. The fixed 1-s design cannot identify a recovery time constant or distinguish adaptation from other causes of depression.

## Discussion

We find no evidence of predictive anticipatory activity in the largest public in-vitro stimulation dataset we found, and the timing of the paradigm explains the apparent pre-stimulus structure we examined. The null is bounded by equivalence tests, was specified in advance and tested on held-out experiments, agrees with the pre-specified sensitivity analyses, and can be reproduced from the public files with hash-verified code. It does not depend on the significance threshold of any single test.

Four limits qualify the conclusion. The first is sensitivity. The pre-specified biological positive control, growth of decoding over the repeated sequence, was itself null, and the closed-loop dataset the plan named as a second positive control (DANDI 001132, human hippocampal slices in which epileptiform activity triggers optogenetic stimulation) has no stimulation tables. The bound therefore rests on synthetic injection of multiplicative rate changes in 250-ms spike counts. We did not show that this preparation can express pre-stimulus label information in any form, and a signal carried by spike timing or spatial pattern would be invisible to both questions. The second is maturity. The rat cultures were recorded at DIV 8-35 over the deposit; session 1 of the 13 held-out Q1 experiments fell at DIV 8-24, derived from the session recording dates (the deposit’s per-subject DIV field disagrees with those dates for one culture and was not used directly). In the same preparation^15^, even basic deviance detection, a larger response to a rare stimulus than to a frequent one, is absent at DIV 10 and emerges only from about DIV 15, so the youngest sessions predate the stage at which prediction-related responses appear. Cultured neurons^16^ and mouse cortical organoids^17^ can learn when embedded in a closed-loop task, whereas open-loop stimulation protocols in dissociated cortical cultures have mostly produced changes no larger than those that occur spontaneously^18^. Repeated stimulation at short intervals also suppresses response magnitude in this preparation^18^; Q2 shows only that 100 repetitions of a sequence in this paradigm produced no detectable pre-stimulus information about the next stimulus, not that the cultures cannot learn. The third is design. Event times in the hidden-source experiment are fixed to a 1-s grid, so the experiment tests anticipation of unpredictable content only, not of unpredictable timing. In the hidden-source experiment the draw is unpredictable only on first exposure, and sessions within an experiment are not independent replicates; position-level inference and the nesting of experiments within cultures account for this. The fourth is one pre-specified interpretation control, growth in decoding of the label five trials ahead, which remains unexplained although it did not replicate on the independent development split (Results).

These results also specify what a decisive in-vitro test needs. Event times should be drawn from a flat-hazard distribution, in which the chance of an event does not change with the time already elapsed, so that no temporal expectation can form. The design needs at least two contrastable stimulus categories, interleaved no-stimulus catch trials, per-electrode spatial tests, and enough data for power at the human-reported effect sizes^3^: several thousand trials per culture and ten or more cultures. The present analyses supply the effect-size bounds and internal timing control for planning it. Until it is run, the claim should stay with what was measured: in these rat cortical cultures, at these ages, anticipation of unpredictable events was not detected at the stated sensitivity.

## Resource availability

### Lead contact

Further information and requests should be directed to and will be fulfilled by the Lead Contact, Cedric Cannard.

### Materials availability

This study did not generate new materials.

### Data and code availability

- This paper analyzes existing, publicly available data from the DANDI Archive: the hidden-source inference recordings of rat cortical cultures^19^ (dandiset 001611, version 0.260611.0634; DOI 10.48324/dandi.001611/0.260611.0634) at https://dandiarchive.org/dandiset/001611/.
- The analysis code, the frozen specification and its parameter file, the held-out split and the saved run outputs are public at https://github.com/amisepa/cortical-anticipation-001611 (GPL-3.0) and will be archived with a DOI at acceptance. The run’s summary file records the SHA-256 of every frozen file, and the OSF registration summary records those of the three frozen plan documents; a script in the repository recomputes them, and all match (last checked 2026-11-09; STAR Methods, Additional resources).
- Any additional information required to reanalyze the data reported in this paper is available from the lead contact upon request.

## Acknowledgments

The authors thank the team behind DANDI dandiset 001611 for making these data public. This work received no specific funding.

## Author contributions

C.C.: Conceptualization, Methodology, Software, Formal analysis, Investigation, Data curation, Writing (original draft; review and editing), Visualization. W.R.: Conceptualization, Methodology, Software, Formal analysis, Validation, Investigation, Data curation, Writing (original draft; review and editing).

## Declaration of generative AI and AI-assisted technologies in the manuscript preparation process

During the preparation of this work, the authors used Claude Opus 5.5 and Claude Fable 5.1 and 5.2 (Anthropic) to write, run and debug analysis code under the authors’ supervision, to draft the analysis plan, checkpoint reports and decisions log, to search open data archives, to draft and edit text, check references, and check and document the public code repository; Gemini (Google) to cross-check the manuscript; and Grok (xAI) for an early review of the analysis plan. The authors reviewed and edited all output and take full responsibility for the content of the published article.

## STAR Methods

### Key resources table

| REAGENT or RESOURCE | SOURCE | IDENTIFIER |
|---|---|---|
| **Deposited data** |  |  |
| Hidden-source inference HD-MEA deposit (DANDI 001611) | Mayama et al.<sup>19</sup> | https://dandiarchive.org/dandiset/001611/ (version 0.260611.0634; DOI 10.48324/dandi.001611/0.260611.0634, CC-BY-4.0) |
| Analysis code and frozen specification (Python/R) | This paper | https://github.com/amisepa/cortical-anticipation-001611 (archival DOI at acceptance) |
| **Software and algorithms** |  |  |
| Python | Python Software Foundation<sup>20</sup> | version 3.11.18; https://www.python.org |
| scikit-learn | Pedregosa et al.<sup>21</sup> | version 1.9.1; https://scikit-learn.org |
| NumPy | Harris et al.<sup>22</sup> | version 2.4.6; https://numpy.org |
| SciPy | Virtanen et al.<sup>23</sup> | version 1.17.1; https://scipy.org |
| pandas | McKinney<sup>24</sup> | version 3.1.0; https://pandas.pydata.org |
| pynwb | Rübel et al.<sup>25</sup> | version 4.2.0; https://pynwb.readthedocs.io |
| h5py | h5py developers<sup>26</sup> | version 3.16.0; https://www.h5py.org |
| R | R Core Team<sup>27</sup> | version 4.5.3; https://www.r-project.org |
| metafor | Viechtbauer<sup>28</sup> | version 5.0-1; https://www.metafor-project.org |

### Experimental model and study participant details

All recordings are previously published public deposits (see Data and code availability); this study is a computational re-analysis and involved no new experiments, subject-derived materials, or human participants. Dandiset 001611 (version 0.260611.0634) contains twelve dissociated rat cortical cultures (Rattus norvegicus, Wistar strain, plated from embryonic day 18 cortical tissue) on high-density multi-electrode arrays (MaxOne geometry, up to 1,024 recording channels), performing a hidden-source inference task. We downloaded all 2,700 NWB session files and verified each against its deposited SHA-256. The files group into 27 experiments of 100 consecutive sessions each (100 to 600 sessions per culture), every session holding 256 trials: 691,200 trials in total. The deposit contains channel-level threshold detections, not raw traces, and stores event times in milliseconds. Culture age at each session was taken from the recording dates (days in vitro, DIV, 8-35 over the deposit; 20 of 27 experiments start at DIV 15 or later; supplement Table S4); the DIV field of the subject description is inconsistent for one subject and was not used directly. Each trial draws two hidden binary sources, s¹ and s², and delivers an observation as a 32-electrode stimulation volley in which electrodes 1-16 encode s¹ and 17-32 encode s² with asymmetric per-electrode reliability (0.75 / 0.25); a (0,0) draw delivers no stimulation. These no-observation trials are one of the four categories and are coded as such throughout (about 25 percent of trials; 19.9-30.1 percent per experiment).

### Method details

#### Trial timeline and the pre-stimulus window

The deposit has no stimulation table, so stimulation events were reconstructed label-free from mass-synchronous artifact detections: a volley is at least three 0.25-ms bins, each with at least 10% of channels detecting, within -16 to +12 ms of the recorded trial time. Electrical stimulation on multi-electrode arrays produces a transient artifact that reaches many electrodes at once, can saturate the amplifiers near the stimulation sites, and makes threshold detection unreliable for milliseconds after each pulse^29,30^; in this deposit each volley therefore appears as near-simultaneous detections on many channels. Every stimulated trial has exactly one volley, lasting from about -5 ms to +7 ms around its registration time (onset as early as -13 ms in one culture), and no other stimulation occurs anywhere in a session. Trials fall on an exact 1.000-s grid (per-session Theil-Sen fit; residuals within ±0.025 ms in the audited experiment C03_E02 and at most 0.78 ms in any of the 2,700 sessions). Earlier volleys therefore sit at exactly -1, -2, -3 s relative to a trial’s own volley. No-observation trials have no volley (0 mismatches between the label and the detected volleys in 332,800 development-split trials; three held-out sessions contain one more detected volley than the labels imply, an aggregate check not inspected at trial level). The recorded registration timestamps carry a label-dependent jitter of one sample (50 µs): no-observation trials are registered about 0.05 ms earlier. We therefore anchor every trial to t₀(n), the fitted 1-s grid plus the experiment’s median detected volley onset, and never use the recorded start times or inter-trial intervals as covariates. The pre-stimulus window is W(n) = [t₀(n) - 265 ms, t₀(n) - 15 ms], a fixed 250-ms window for every experiment and trial. It is stimulation-free by construction and checked per trial: a trial whose window overlaps any detected volley’s blanking span (onset - 4 ms to offset + 3 ms) is dropped (1 of 691,200 trials, in a session no confirmatory analysis uses). It is not free of the previous trial’s after-effects: at a 1-s interval no window is. Evoked activity decays over several hundred milliseconds, and the window rate after a volley, as a ratio to the rate after a no-observation trial, ranges from 0.07 to 3.4 across held-out experiments. This carryover is removed statistically rather than by excluding windows, which with a fixed sequence would select trial positions by their previous label. The first two trials of each session are dropped (no history), leaving up to 254 trials per session. Channels with an artifact-free rate of at least 0.01 Hz inside the windows over the whole experiment are included (label-free; 600-974 channels per held-out experiment). Sessions are numbered 1 to 100 in this text; the split file and the code count from 0, so their sessions 0-49 are sessions 1 to 50.

Question 1: anticipation of an unpredictable draw (primary). In session 1 of each experiment, does activity in W(n) carry information about s¹(n) and s²(n) beyond trial history? Features: log1p of the total window count and of the counts in three spatial pools defined from array geometry (channels within 100 µm of a source-1 electrode, of a source-2 electrode, and all others). History H(n), 11 covariates: s¹ and s² of trials n-1 and n-2, the number of electrodes stimulated in each half of the previous trial’s volley (its observation, o(n-1)), the log1p evoked counts 25-300 ms after t₀(n-1) (total and three pools; spontaneous counts in that interval for no-observation trials) and trial position. Features are residualized on H(n) by ridge regression (alpha = 1) fitted inside each training fold only (cross-fitted). Two L2-regularized logistic regression decoders (inverse regularization strength C = 1), one per source, on standardized residualized features. Cross-validation uses 5 contiguous blocks of trial positions with a 5-position gap on each side of the test block. The per-experiment statistic is the mean over the two sources of the mean within-fold AUC.

Question 2: learned expectation (secondary confirmatory). Across the 100 sessions of an experiment, does history-residualized pre-stimulus decodability grow as the sequence repeats? Features: log1p per-channel window counts. H(n) adds the session index, the previous session’s mean log1p total window count, the cumulative number of stimulation pulses, source frequencies over the last 10 and 50 trials and 5 in-fold principal components of the per-channel evoked response r(n-1). Decoder: L2-regularized logistic regression, C = 0.01 (chosen by injection recovery on the development split, which was flat from C = 10⁻⁴ to 10⁻²). Sessions are grouped into 10 blocks of 10; within each block the same position-aligned 5-fold scheme is used, so a test position never appears in training. The per-experiment statistic is G = 9 × the OLS slope of block AUC on block index, the fitted change from block 1 (sessions 1-10) to block 10 (sessions 91-100). Secondary contrast: block 10 minus block 1. Q2 also serves as the biological positive control for Q1: if cultures come to expect the repeated sequence, residualized decodability should grow. It is the only test of learned expectation in these data and is never interpreted as anomalous anticipation.

#### Pre-task timing control

For each of the 2,700 sessions, population spike times were binned at 10 ms from time 0 to the last bin boundary at or before 16 ms before the first trial (pre-task) and for an equal duration from the first trial (task). Ending the pre-task segment 16-26 ms before registration keeps the leading edge of the first volley, which precedes registration by about 5 ms, out of its last bin. Normalized autocorrelations were calculated at lags 0.10-2.00 s after subtracting each segment’s mean; each lagged covariance was normalized by the segment variance. Volley detections were kept in the task segment, whose autocorrelation (0.36) therefore reflects the stimulation clock itself; with the 10-ms bins within -16 to +12 ms of each trial time masked it was -0.03. Autocorrelations were averaged within each culture and then across the 12 cultures. All sessions had at least 20 pre-task spikes (median 234), so none were excluded. This analysis was descriptive and outside the registered plan.

#### Response recovery and within-session depression

These descriptive analyses used all 2,700 sessions and the deposited spike detections. For each trial, spikes were binned in 50-ms bins from 0 to 1000 ms after its recorded registration time. Rates were normalized by the number of sorted units in the file, the number of trials and the bin width. The evoked rate was measured at 100-200 ms and the late rate at 600-900 ms. No-observation trials supplied the matched late-activity comparison. Within each session, evoked counts at 100-300 ms were calculated separately for the three stimulated categories and for trials 1-128 and 129-256. For each experiment, the second-half/first-half ratio was the geometric mean of the within-category session ratios, and a category entered that ratio only if its mean evoked response was above zero in both trial halves, averaged over the first ten and the last ten sessions. This analysis measures response magnitude and is not a test of category selectivity or recovery at other intervals.

#### Development and held-out split

Experiments were split before any outcome analysis (split file SHA-256 2d792f9e…65ef9): in each multi-experiment culture, half the experiments (rounded up) were held out; in the four single-experiment cultures, sessions 1-50 and 51-100 were assigned to the two splits at random. The analysis unit is an experiment, or one half of a culture recorded in a single experiment. All development used the development split (15 units), with permuted labels until the pipeline was fixed. All three real-label controls ran after the primary run (2026-10-28, 21:57-23:03 PDT): the lag profile and the timing-anchor check on development data, as scheduled before the held-out run, and the post-stimulus check on held-out data. The timing-anchor check was repeated on the held-out Q1 experiments on 2026-10-29. The held-out split was used before the run only for label-free integrity checks and labels-only sequence checks. The data were public and had been analyzed by one author in earlier analyses, including the held-out experiments, before the plan was fixed; the plan is pre-analysis for this pipeline on the held-out split, not pre-data. Q1 confirmatory set: session 1 of the 12 held-out experiments of multi-experiment cultures plus the one single-experiment culture whose held-out half contains session 1 (13 experiments in 9 cultures). Q2 confirmatory set: the 12 held-out experiments of multi-experiment cultures (12 experiments in 8 cultures); single-experiment cultures are excluded because both halves share one sequence and one culture. Five Q1 experiments start before DIV 15 (C01_E02, C04_E01, C05_E02, C07_E01, C09_E01), so the age sensitivity has 8 experiments in 6 cultures (Q2: 7 in 5). Power at the realized design, from injection on the development split, is in Table S2.

Pipeline (frozen spec sections in parentheses; every frozen file is pinned by SHA-256 in the run summary, and the three plan documents also in the OSF registration summary). Preprocessing (spec 2-3): all 2,700 NWB session files were verified against their deposited SHA-256 hashes, and t₀(n) was estimated per session as described above; the recorded start_time was never used directly because its jitter is label-dependent. Artifact handling (spec 4): the span from volley onset - 4 ms to offset + 3 ms was treated as artifact, but no detection was deleted; the span decides trial inclusion, and the post-stimulus interval (25-300 ms after t₀) starts after it. Trial and channel inclusion (spec 3) were label-free, as described above, and preprocessing removed no further analyzed trial. The specification also excludes channels sitting on a stimulation electrode; no recording channel does (the nearest lies 17.5 µm from a stimulation electrode in every session), so this rule removed no channel. Features, history covariates, decoders and position-aligned cross-validation (spec 4-6) are given under Questions 1 and 2; features were residualized by ridge regression on standardized covariates, fitted within each cross-validation training fold only (cross-fitted residualization, spec 5). The sequence-shift null, pooling and the randomization test (spec 6.5) and equivalence testing and Bayes factors (spec 7) are described under Quantification and statistical analysis. The complete pipeline, with the frozen specification and the saved run outputs, is public in the analysis repository^31^.

#### Freeze and run timeline

The analysis plan (the specification, its parameter file, the pipeline code and the two entry scripts) was frozen at git tag prereg-v1 (commit 71e5018) on 2026-10-28 at 16:19:20 PDT; the tag pins every frozen file by hash, and its time is set by the local machine. The held-out run started 19:06:44 PDT from a checkout that a guard script verified to be identical to the tag, with the hashes of the split file, the environment lock and the pre-run gate results matching, and ended 20:08:13 PDT; no frozen file changed. The Open Science Framework registration (https://osf.io/nxsph/) was created at 19:51 PDT, during the run and before any per-experiment output was written (19:58:46), and holds the SHA-256 hashes of the three frozen plan documents. Three pre-specified controls ran after the primary analysis (21:57-23:03 PDT, with operationalizations committed before running), and two later uses of held-out data that cannot change an outcome (a repeat of the timing-anchor check on 2026-10-29 and the exploratory per-fold recomputation of C01_E02) are departures from the specification’s runs-once wording, stated as such. The plan also lists further exploratory analyses that are not reported here: a re-run of our earlier exploratory analysis and its other tests (families S1 and S2), secondary targets (S3), cross-dataset comparisons (S4), run-length models and a per-session Q2 curve (S5), and a cross-preparation comparison of spontaneous activity (specification section 13). Of these, one was run, before the freeze: a leakage diagnostic on a sister dataset (Isomura et al.; specification 4c), which decoded the source of trial n from the evoked response to trial n-1 and was null (AUC 0.491 against a null mean of 0.500, where post-stimulus decoding on the same data reached 0.996). The others were not run. None can change the outcome for Q1 or Q2. Three inputs that determine the unit set and the age subset (the DIV table, the session index and the unit list) were not hashed by the guard; their hashes were recorded afterwards, the two tracked files are unchanged since the tag, and the session index (not under version control) predates it. The full timestamped timeline, including the fresh-unzip SHA-256 re-verification of 2026-11-02, is in Table S6. C.C. reproduced every held-out per-experiment AUC, shift-null mean and shift-null variance (all 13 Q1 and 12 Q2 experiments) from the public NWB files with the frozen code of the run snapshot; all agree with the snapshot’s tables at their printed precision, and the pooled estimates and standard errors reproduce the run summary’s printed values.

#### Software

Versions are listed in the Key resources table (Python 3.11.18 with scikit-learn 1.9.1, numpy 2.4.6, scipy 1.17.1, pandas 3.1.0, pynwb 4.2.0 and h5py 3.16.0; R 4.5.3 with metafor 5.0-1 for pooling). Bayes factors use a normal likelihood on the pooled estimate and its standard error, two-sided, with a Cauchy(0, SESOI) prior and robustness at 0.5× and 2× the scale (frozen spec, Section 7). The environment lock file is hashed in the run summary (SHA-256 c8364adf…3a2). Global seed 20261026, with per-task seeds derived deterministically. The analysis code and the frozen specification are available at https://github.com/amisepa/cortical-anticipation-001611.

### Quantification and statistical analysis

#### Inference at the position level

Sessions of one experiment repeat the same label sequence, so the unit of replication for any label-dependent statistic is the trial position (254 per experiment), not the trial. No trial-level bootstrap or per-trial test is used. Sequence-shift null: the 256-trial label sequence is circularly shifted by every k from 16 to 240 (225 shifts), the same shift in every session, and the identical pipeline is rerun with the shifted sequence as the target only (history unshifted, because it describes the physical past of the window). The per-experiment effect is the observed statistic minus the mean of its null (ΔAUC for Q1, ΔG for Q2), with sampling variance equal to the null variance. Pooling: a three-level random-effects model with known sampling variances, experiments nested in cultures (REML; R metafor rma.mv, random = ∼1 | culture/experiment), t-distribution with df = cultures - 1. Primary p-value: a randomization test that draws one shift-null value per experiment, refits the same model, and repeats 10,000 times. A sequence-swap null (each experiment’s windows paired with the position-aligned labels of every other experiment’s sequence in the same split) is reported as a check that the shift null is centered. Forest plots show each experiment’s estimate with the 95% interval of its own shift null.

#### Decision rules and equivalence

Smallest effect sizes of interest (SESOI) were fixed before the run. For Q1, the smallest effect of interest is ΔAUC 0.059, the conversion of the pooled human-reported effect (d ≈ 0.21; ΔAUC = Φ(d/√2) - 0.5), with a secondary bound of ±0.03 reported as such. For Q2, it is 0.02 (growth in AUC from block 1 to block 10). For each question: equivalence by two one-sided tests at ±the smallest effect of interest (alpha = 0.05, equivalent to the 90% CI lying inside the bounds), the randomization p, and a Bayes factor with a Cauchy(0, SESOI) prior with robustness at 0.5× and 2× the scale. Each question had three possible outcomes: B, a candidate effect, if the 95% CI excludes 0, the randomization p is below 0.05 and every control passes, whatever the size; A, a bounded null, if not B and equivalence holds; and C, inconclusive, otherwise. The realized 90% CI is reported as the achieved bound. Q1 and Q2 are separate families of one test each, with no multiplicity correction between them. Sensitivity analyses are reported, not tested: a 360-ms window with the same end; experiments at DIV ≥ 15 at session 1; unresidualized, rank-transformed and all-channel features (Q1).

#### Controls

All controls were specified before the run; the qualitative pass criteria of the post-stimulus, lag-profile and timing-anchor controls were operationalized after the primary result was known, and the real-label development controls had been scheduled to run before the held-out run. Two checks had to pass before the held-out run: (a) on the development split, after residualization, windows cannot decode the previous trial’s label s(n-1) (pooled excess over permuted-label runs, with a 95% CI upper bound below 0.03 for Q1 and 0.02 for Q2 and a lower bound at or below 0); (b) on the labels of all 27 sequences, not only the development split, a position-level randomness audit (16 Monte Carlo tests per sequence against independent fair coin flips, Holm over 432 tests; 18 global tests, Bonferroni). Other controls: injection of Poisson spikes that multiply a channel’s expected window count by (1 + δ)^(s¹+s²) on real data with permuted labels, which gives the false-positive rate, the detectable effect at 80% power and, inside the confirmatory run, each experiment’s injected sensitivity (Q1: δ = 0.05 on all included channels; Q2: a ramp to δ = 0.1 over the 100 sessions); post-stimulus decoding of s(n+k), k = -2…+2, from counts 25-300 ms after t₀ (the pre-specified rule says “well above 0.5”; its operationalization, that the peak must be at k = 0 in every experiment and block, was fixed when the control was run); a lag profile decoding s(n+k) from W(n) for k = -3…+10 on real development labels (targets 5-10 trials ahead must be null); a timing-anchor check that re-runs the Q1 decoder with windows anchored on the recorded start times instead of the grid (any difference raises a flag); a position-lookup check that naive by-session cross-validation decodes permuted fixed-sequence labels and position-aligned cross-validation does not; preprocessing balance across categories at the position level; and three Q2 interpretation controls: the per-block unresidualized AUC for s(n-1) (carryover trajectory), growth G for the targets s(n+5) and s(n+10) against the same shift null (flagged, meaning investigate, if either 95% CI excludes 0), and the Q2 swap null. Table S5 summarizes the controls and their status.

### Additional resources

#### Pre-analysis plan registration

Open Science Framework registration of the frozen analysis plan, created 2026-10-28 at 19:51 PDT during the confirmatory run (started 19:06:44 PDT) and before any output was written (19:58:46 PDT), embargoed during peer review and made public at acceptance. Description: https://osf.io/nxsph/ (a view-only link provides access to reviewers on request). The registration holds the SHA-256 hashes and git blob IDs of the three frozen plan documents; the frozen code and the saved run outputs are in the public code repository (Data and code availability).

## Supplemental Information

### Methods S1. Supplementary analyses and data audits

#### Methods S1.1. The confirmatory window, the earlier exploratory window and the artifact rule

Our earlier analysis used the 3.45–3.10 s window before each trial’s registration time as the confirmatory window. On the 1-s trial grid (Fig. 1) that window lies 0.55–0.90 s after the volley of trial n−4 and ends 95 ms before the volley of trial n−3. Its spike counts therefore test whether activity before trial n−3 carries the category of trial n, with three stimulated trials in between: a lag-3 question. Its first 160 ms lie in the response tail of trial n−4. The “engine slots” at about 2.45, 0.90, 0.50 and 0.05 s before registration correspond to the volley of trial n (−0.005 s; the volley sits about 5 ms before registration, and the earlier analysis’s 100-ms bins place it at about 0.05 s) and to evoked-response phases of the volleys at −1.005 and −3.005 s (0.90 s before registration is about 100 ms after volley n−1, near the evoked peak; 0.50 and 2.45 s fall in the response tails). No stimulation event occurs at 2.45, 0.90 or 0.50 s in any of the 101 audited sessions.

Our lag profile puts the same question in the corrected framework. Decoding the label of trial n+k from the window of trial n on real development-split labels (14 Q1 units; 11 Q2 units; history removed for k ≥ 0) gives, at k = +3, ΔAUC −0.021 (95% CI −0.048 to +0.006) for session 1 and +0.000 (−0.003 to +0.003) for the Q2 block AUC. The full profile is in Table S1. The k = +3 point above is the corrected-framework analog of the earlier window; the earlier analysis’s own estimates are not reported here.

#### Methods S1.2. Data notes: what the deposited files contain, and what they do not

1.**Fixed sequences.** Each of the 27 experiments uses one 256-trial (s¹, s²) sequence in all 100 sessions; identity verified in every session; no reuse across experiments, including circular shifts and source swaps or flips (maximum shifted or swapped agreement 0.60). Simultaneous experiments on two chips use different sequences.
2. **One volley per trial.** Volleys at −5 to +7 ms around registration (onset as early as −13 ms in one culture), on a 1.000-s grid (residuals ±0.025 ms in the audited experiment C03_E02; at most 0.78 ms across the 2,700 sessions); no other stimulation; no-observation trials have none. The 244-s inter-session rest is not in the deposit.
3. **Stimulation artifacts.** In the audited experiment (C03_E02, 100 sessions, plus one partner session), 80–96% of task-period detections per session fall within the volley spans (pooled 88.9%; 84.5–90.4% depending on the span definition). Artifact-inclusive rate 6.4 detections per second per channel; artifact-free 0.72 (0.24–1.46 across sessions); spontaneous rate before the first trial 0.026. The 4.8 per second per channel of our earlier analysis is an artifact-inclusive figure.
4. **Carryover.** The window rate after a volley divided by the rate after a no-observation trial ranges from 0.07 to 3.4 across held-out experiments (0.15 to 1.99 on dev): some cultures are suppressed after a volley, others reverberate.
5. **Housekeeping.** Times are stored in milliseconds. No-observation trials are registered one sample (50 µs) earlier than stimulated trials, a label-dependent jitter; windows are anchored on the fitted grid for this reason, and the recorded inter-trial interval must not be used as a covariate. The subject DIV field is inconsistent for one subject; age is derived from recording dates (Tokyo calendar days).
6. **Channel-level detections only.** The deposit holds threshold crossings per channel, not raw traces, so a spike-detection census cannot be applied to it on the same terms as to raw datasets.

#### Methods S1.3. The flagged 5-trials-ahead control

##### Registered control

Growth G for the targets s(n+5) and s(n+10), computed exactly as the Q2 statistic against the same shift null, flagged if either pooled 95% CI excludes 0. Held-out: k = 10 +0.0005 (−0.0075 to +0.0086); **k = 5 +0.0105 (SE 0.0036; +0.0019 to +0.0191; REML p 0.023; randomization p 0.0025; positive in 10 of 12 experiments)**. The L360 sensitivity run gave +0.0038 (−0.0053 to +0.0130) at k = 5.

**Plan, fixed before running** (committed 2026-10-29 11:24:46 PDT; run 11:27–14:02, once, no changes; development data and saved held-out outputs only): (a) the registered randomization p for the held-out value, from the saved nulls; (b) the same statistic on the development split, the independent replication; (c) labels-only common-mean auto- and cross-correlations of the 27 sequences at lags 1–12 against 10,000 within-sequence permutations; (d) the growth profile over k = 1…10 on dev, leave-one-out and per-block views. Rules: shared lag-5 structure in (c) → a property of the generator; (b) null and (c) null → chance finding, reported with the multiplicity count; (b) replicates and (c) null → unresolved anomaly; a smooth profile across k in (d) → the Q2 bound quoted as ±0.03 in effect.

##### Results

(b) Dev (11 experiments, 8 cultures): ΔG₅ +0.0035 (−0.0062 to +0.0132), randomization p 0.33, positive in 7 of 11: null. ΔG₁₀ −0.0013 (−0.0116 to +0.0090). (c) No lag-5 test is significant (common means −0.027, −0.016, −0.002, −0.022 for auto s¹, auto s², cross s¹→s², cross s²→s¹; smallest Holm p 0.21 over the 4 tests; 0.12 under the uncentered formula); the lag-5 means are negative. (a) Held-out ΔG₅ randomization p 0.0025; Holm over the 2 registered growth controls 0.005; Holm and Bonferroni over the 14-test family (2 growth controls + 12 lag tests at k = 5…10 for Q1 and Q2) 0.035; all 13 other tests have Holm p 1.0. (d) On dev every pooled ΔG_k for k = 1…10 has a CI including 0 (range −0.003 to +0.007; k = 5 mid-range); the mean over k ≠ 5 is +0.0024 (−0.0002 to +0.0051) and the contrast G₅ − Ḡ_other is +0.0010 (−0.0080 to +0.0101): neither smooth nor k-specific. Held-out leave-one-experiment-out estimates stay above 0 (+0.0094 to +0.0126); leaving out culture C04 (3 experiments) gives +0.0081 (−0.0015 to +0.0176). **Rule 2 fires: chance finding. Rule 4 does not fire: the Q2 bound stays ±0.02.** The flag did not replicate on the development split, and the sequences show no shared lag-5 structure; it is classified as chance under the pre-stated rule but remains unexplained. The development estimate contains the held-out value, so non-replication is weak evidence, and the flag stays nominally significant after Holm correction over the 14-test family. A lag-8 path that rule 1 did not test is described in Methods S1.4. Nothing further was run (stop rule).

#### Methods S1.4. Within-sequence randomness audit

Of the 48 common-mean statistics in Methods S1.3(c), one has a Holm-adjusted p below 0.05: the circular autocorrelation of s¹ at lag 8, common mean +0.037 over the 27 sequences (permutation null mean −0.004, SD 0.012; 20 of 27 sequences positive). Two formulas apply, because a permuted circular autocorrelation has expectation −1/255 rather than 0: under the formula pre-specified in the committed script, centered on the null mean, p = 0.0008 and Holm over 48 = 0.038; under the plan’s literal uncentered formula, p = 0.0041 and Holm over 48 = 0.20. The lag-6 value is +0.027 (p 0.0098, Holm 0.46). This is a property of the stimulation sequences, not of the recordings; it is outside every rule and was not followed up (stop rule). It corresponds to s¹(n) and s¹(n+8) agreeing on about 52% rather than 50% of trials; whether it reflects the generator or chance among 48 tests cannot be settled from 27 sequences. The per-sequence audit (gate (b)) has little power against such shared structure, so its pass does not establish independence at the pooled level. The primary targets are not exposed: the common autocorrelations at lags 3 and 4, where residual carryover could act, are null (s¹ lag 3 +0.007, p 0.36; lag 4 +0.006, p 0.42; s² p ≥ 0.48). For the k = 5 control, a lag-8 dependence links s(n+5) to s(n−3), whose carryover remains detectable in the windows (Table S1, Q2 k = −3: +0.007, CI +0.001 to +0.014); this path has a consistent sign across sequences, was not covered by the lag-5 test in rule 1, and was not tested. A rough post-hoc linear bound, not a project output: the path would contribute about 0.04 × 0.007 ≈ 0.0003 to ΔG₅, roughly 30 times smaller than the flagged +0.0105, so it is unlikely to explain the flag; nothing was run to check this. It is listed for the follow-up paper.

#### Methods S1.5. Controls checklist

The controls and their status are listed in Table S5.

#### Methods S1.6. Freeze and run timeline

The freeze and run timeline is listed in Table S6.

#### Methods S1.7. Power at the realized design

Power at the realized design is summarized in Table S2.

#### Methods S1.8. Per-experiment results

ΔAUC and ΔG are the experiment’s statistic minus the mean of its 225-shift null; the per-experiment null SD is about 0.035 (Q1) and 0.011–0.014 (Q2). Injected sensitivity: Q1, ΔAUC at a 5% rate change on all included channels; Q2, ΔG at a ramp to 10%, both against permuted label sequences.

#### Methods S1.9. The Q1 outlier C01_E02

C01_E02 (DIV 13) has the most negative Q1 value, AUC 0.393 against a null mean of 0.497. Its per-fold, per-source AUCs, recomputed with the frozen code (reproducing the saved values exactly), show one position block well below its null (fold 1, source 1: 0.17, below all 225 shifts) and a broad shift below 0.5 in the other folds (9 of 10 fold-source values below their null mean; leaving fold 1 out still gives about 0.42). The shift null is centered near 0.50 in every fold, so the null construction does not produce below-chance values in general. Leaving the experiment out moves the pooled Q1 estimate to about +0.012; the outcome is unchanged. Nothing further was run.

#### Methods S1.10. Exploratory family S5: secondary contrast and pre/post category-mirror check

Two checks were pre-specified as exploratory family S5 and are reported for completeness. (i) Decoding the (0,1) versus (0,0) categories in session 1, in the same framework as Q1: ΔAUC +0.013 (95% CI −0.032 to +0.059, p = 0.52). (ii) Mirroring of the pre-stimulus category contrast onto the post-stimulus contrast (Spearman across experiments): rho = 0.09, sign agreement 8 of 13, shift-null p = 0.77.

## Tables

**Table S1.** Lag profile on real development labels, related to Figure 1 and STAR Methods. ΔAUC against each statistic’s own shift null, with 95% CIs; rows k ≤ −1 are not residualized.

| **k** | **Q1 (session 1, 14 units)** | **Q2 block AUC (11 units)** |
|---|---|---|
| −3 | +0.010 (−0.012 to +0.032) | +0.007 (+0.001 to +0.014) |
| −2 | +0.039 (+0.005 to +0.073) | +0.027 (+0.009 to +0.045) |
| −1 | +0.044 (+0.013 to +0.075) | +0.078 (+0.040 to +0.116) |
| 0 | −0.001 (−0.022 to +0.020) | −0.001 (−0.004 to +0.002) |
| 1 | +0.006 (−0.015 to +0.027) | +0.001 (−0.004 to +0.006) |
| 2 | −0.008 (−0.029 to +0.013) | +0.002 (−0.001 to +0.005) |
| 3 | −0.021 (−0.048 to +0.006) | +0.000 (−0.003 to +0.003) |
| 4 | −0.004 (−0.033 to +0.024) | −0.001 (−0.004 to +0.002) |
| 5 | −0.014 (−0.035 to +0.007) | +0.002 (−0.001 to +0.005) |
| 6 | −0.004 (−0.027 to +0.019) | +0.001 (−0.002 to +0.004) |
| 7 | +0.009 (−0.012 to +0.030) | +0.000 (−0.003 to +0.003) |
| 8 | +0.010 (−0.011 to +0.031) | −0.000 (−0.005 to +0.004) |
| 9 | +0.005 (−0.016 to +0.027) | +0.000 (−0.003 to +0.003) |
| 10 | −0.002 (−0.023 to +0.019) | +0.001 (−0.003 to +0.004) |

Note that the k = 0 row is the development-split level of the Q1 statistic on real labels (not a confirmatory test), and the Q2 rows are block-level AUCs, not the growth statistic.

**Table S2.** Power at the realized design, related to Figures 2 and 3 and STAR Methods.

|  | **Q1, 14 dev units (11 cultures)** | **Q1, 9 cultures (held-out-like)** | **Q2, 11 dev units (8 cultures)** |
|---|---|---|---|
| 80% detection, homogeneous effect | ΔAUC 0.031 | 0.036 | ΔG 0.013 |
| 80% detection, rate-proportional injection, all channels | 0.054 | 0.063 | 0.050 |
| 80% detection, injection on pools A+B / top-8 channels | 0.049 / 0.074 | 0.060 / 0.088 | 0.080 / 0.090 |
| Detection power at the SESOI, homogeneous / all-channel injection | 1.00 / 0.90 | 1.00 / 0.75 | 0.99 / about 0.2 |
| Equivalence power at ±SESOI under the null | 1.00 | 1.00 | 1.00 |
| Equivalence power at ±0.03 (Q1 secondary) | 0.73 | 0.54 | n/a |
| False-positive rate (nominal 0.05) | 0.01–0.03 | 0.01 | 0.00–0.01 |
| Mean pooled SE under the null | 0.0105 | 0.0117 | 0.0041 |

Realized held-out pooled SE: Q1 0.0129, Q2 0.0042. A rate-proportional effect is very unequal across experiments, so a heterogeneous Q2 growth of 0.02–0.03 would most likely have been “inconclusive” rather than detected.

**Table S3.** Per-experiment held-out results, related to Figures 2 and 3.

| **Experiment** | **DIV at session 1** | **Channels** | **Carryover ratio** | **Q1 ΔAUC** | **Q1 injected** | **Q2 ΔG** | **Q2 injected** |
|---|---|---|---|---|---|---|---|
| C01_E02 | 13 | 953 | 1.33 | −0.105 | +0.155 | +0.008 | 0.170 |
| C01_E03 | 15 | 921 | 1.20 | +0.038 | +0.224 | −0.019 | 0.085 |
| C02_E01 (sessions 0–49) | 20 | 960 | 0.19 | +0.028 | +0.207 | n/a | n/a |
| C03_E01 | 24 | 600 | 0.61 | +0.039 | −0.004 | +0.005 | 0.019 |
| C04_E01 | 9 | 974 | 0.82 | +0.041 | +0.221 | −0.004 | 0.048 |
| C04_E02 | 23 | 954 | 0.17 | −0.018 | +0.066 | +0.003 | 0.128 |
| C04_E03 | 24 | 963 | 0.07 | −0.004 | +0.081 | +0.008 | 0.093 |
| C05_E02 | 14 | 940 | 1.08 | +0.003 | +0.133 | +0.018 | 0.171 |
| C07_E01 | 8 | 766 | 1.08 | +0.053 | +0.027 | −0.006 | 0.024 |
| C08_E01 | 16 | 967 | 0.65 | −0.032 | +0.286 | +0.027 | 0.119 |
| C09_E01 | 8 | 944 | 3.39 | −0.063 | +0.123 | −0.010 | 0.063 |
| C11_E01 | 16 | 965 | 0.66 | +0.035 | +0.340 | +0.022 | 0.028 |
| C11_E03 | 23 | 947 | 0.99 | +0.033 | +0.011 | −0.010 | 0.020 |

Session ranges in this table and in Table S4 are numbered from 0, as in the split file; the main text counts sessions from 1.

**Table S4.** Experiments, cultures, age at session 1 and split, related to STAR Methods. Age is derived from the recording date of each experiment’s first session relative to the culture’s first recording and the subject’s DIV at that date (Tokyo calendar days). The subject DIV field agrees with the date-derived value for 26 of 27 experiments; for C09_E02 the field reads 13 where the dates give 15, and the date-derived value is used. 20 of 27 experiments start at DIV 15 or later. Single-experiment cultures (C02, C06, C10, C12) were split by session halves.

| **Experiment** | **Culture (author subject)** | **Session 1 date** | **DIV at session 1** | **Split** |
|---|---|---|---|---|
| C01_E01 | C01 (24846) | 2024-11-20 | 8 | dev |
| C01_E02 | C01 (24846) | 2024-11-25 | 13 | held-out |
| C01_E03 | C01 (24846) | 2024-11-27 | 15 | held-out |
| C02_E01 | C02 (25273) | 2025-04-28 | 20 | sessions 0–49 held-out; 50–99 dev |
| C03_E01 | C03 (25876) | 2024-11-08 | 24 | held-out |
| C03_E02 | C03 (25876) | 2024-11-11 | 27 | dev |
| C04_E01 | C04 (25897) | 2024-10-24 | 9 | held-out |
| C04_E02 | C04 (25897) | 2024-11-07 | 23 | held-out |
| C04_E03 | C04 (25897) | 2024-11-08 | 24 | held-out |
| C04_E04 | C04 (25897) | 2024-11-11 | 27 | dev |
| C04_E05 | C04 (25897) | 2024-11-13 | 29 | dev |
| C04_E06 | C04 (25897) | 2024-11-19 | 35 | dev |
| C05_E01 | C05 (26265) | 2024-12-22 | 13 | dev |
| C05_E02 | C05 (26265) | 2024-12-24 | 14 | held-out |
| C06_E01 | C06 (26271) | 2025-04-01 | 15 | sessions 0–49 dev; 50–99 held-out |
| C07_E01 | C07 (26296) | 2025-02-26 | 8 | held-out |
| C07_E02 | C07 (26296) | 2025-03-05 | 15 | dev |
| C08_E01 | C08 (26303) | 2025-03-19 | 16 | held-out |
| C08_E02 | C08 (26303) | 2025-03-27 | 23 | dev |
| C09_E01 | C09 (26342) | 2024-11-20 | 8 | held-out |
| C09_E02 | C09 (26342) | 2024-11-27 | 15 (field reads 13) | dev |
| C10_E01 | C10 (26349) | 2025-03-21 | 18 | sessions 0–49 dev; 50–99 held-out |
| C11_E01 | C11 (29641) | 2025-03-19 | 16 | held-out |
| C11_E02 | C11 (29641) | 2025-03-21 | 18 | dev |
| C11_E03 | C11 (29641) | 2025-03-26 | 23 | held-out |
| C11_E04 | C11 (29641) | 2025-03-31 | 27 | dev |
| C12_E01 | C12 (29713) | 2025-04-25 | 17 | sessions 0–49 dev; 50–99 held-out |

Session ranges are numbered from 0, as in the split file; the main text counts sessions from 1. Source: results/B/B1/div_check.csv and data/manifests/split.csv in the public code repository (both unchanged since tag prereg-v1).

**Table S5.** Controls checklist, related to Figure 1 and STAR Methods.

| # | **Control** | **Result** |
|---|---|---|
| 8.1 | Injection: monotone recovery; false-positive rate at δ = 0 | PASS (dev, permuted labels). FPR 0.00–0.03 |
| 8.2 | Post-stimulus decoding peaks at lag 0 | PASS. Dev: 15/15 units, AUC 0.70–0.98. Held-out: 32/32 tests, AUC 0.61–0.98 (no numeric threshold was fixed before the run; AUC ≥ 0.60 stated post hoc for future runs) |
| 8.3 | Lag profile: k ≥ 5 null, real dev labels | PASS. No k in 5…10 with a CI excluding 0 (Q1 or Q2); carryover peak at k ≤ −1 as expected (Table S1) |
| 8.4 | Sequence-shift null, 225 shifts | Computed as registered |
| 8.5 | Sequence-swap null centered on the shift null | PASS. Swap − shift mean: Q1 −0.0022 (SD 0.013), Q2 +0.0002 (SD 0.003) |
| 8.6 | Randomness audit, position level, 27 sequences | PASS (gate b). Min Holm p 0.80 |
| 8.7 | Closed-loop positive control (001132) | NOT AVAILABLE: no stimulation tables in the deposit |
| 8.8 | Raster review of any effect | NOT APPLICABLE: no effect at the pooled level |
| 8.9 | Preprocessing balance across categories | PASS (trivially): one label-free trial drop, outside every analyzed session |
| 8.10 | Power at the realized design | Table S2 (Methods S1.7) |
| 8.11 | Timing anchor (start_time vs grid), real labels | FLAG under the literal rule; the known one-sample registration jitter. Dev: 32 counts changed in 10 of 14 units, max AUC change 0.0027 (0.08 null SD). Held-out: 26 counts in 6 of 13 units, max 0.00065 (0.017 null SD); the grid version reproduces the saved AUC in all 13 |
| 8.12 | Position-lookup leak demonstrated | PASS (code sanity) |
| 8.13 | Decay and carryover (descriptive) | Carryover ratio 0.07–3.39 held-out, 0.15–1.99 dev |
| 8.14 | Hard gates (a) and (b) | PASS, before the run. (a) Q1 excess +0.002 (−0.012 to +0.016), Q2 +0.0005 (−0.004 to +0.005). (b) min Holm p 0.80, 0 structural mismatches |
| 8.15 | Q2 interpretation controls | (i) carryover AUC 0.574 → 0.612 by block 6; (ii) k = 10 null, **k = 5 flagged**, resolved as a chance finding (Methods S1.3); (iii) swap null centered |

**Table S6.** Freeze and run timeline, related to STAR Methods.

| Time | Event | Evidence |
|---|---|---|
| 16:19:20 | Tag prereg-v1 on commit 71e5018 (primary freeze evidence) | git tag object and commit time |
| 19:06:44 | Held-out run started; guard verified frozen files = tag, split and lock hashes, gates passed | run log; summary.json |
| 19:15 | GitHub push of main and the tag | reflog; session log |
| 19:51 | OSF registration created (https://osf.io/nxsph/; SHA-256 and blob IDs of the three frozen files) | OSF; session log |
| 19:58:46 | Per-experiment outputs written | file times |
| 19:58:48 | Per-experiment file (units.csv) accessed by an unidentified process (consistent with OS indexing) | file access time |
| 20:08:13 | Primary run ended (exit 0); L360 sensitivity run 20:08–21:13 | run logs |
| 20:08:41 | Primary run log read once, for its exit status only | session log |
| 21:13:29 | First time a held-out result was displayed (read-only extraction of summary.json) | session log; file access time |
| 21:14:52 | Both run logs copied into results/ (their birth times). Stdout had been captured from launch in the session's scratch directory, in files born 19:06:44 and 20:08:13 and last modified at each run's end (20:08:13, 21:13:02), byte-identical to the copies; committed at 21:15:05 (5d6875b) | file times; commit |
| 21:57–23:03 | Registered controls 8.2 (held-out), 8.3 and 8.11 (dev) run after the primary analysis; operationalizations committed 17 s before the first output | commit d182dde; file times |
| 29 Oct 11:24:46 | k = 5 investigation plan committed; script 11:27:46; run 11:27:52–14:02:16 | commits 98c4bfc, 5cd675e; run log with commit and hashes |
| 29 Oct 14:02–14:06 | 8.11 on the 13 held-out Q1 units; per-fold look at C01_E02 | commit 78c0552; run logs |

Unknown from the logs: whether anyone viewed held-out output outside the logged sessions; who or what accessed the per-experiment file two seconds after it was written (consistent with OS indexing); the exact second of the tag push (19:14:55–19:15:11). The local tag time (16:19:20) is set by the local machine; the original analysis repository on GitHub is private and the OSF registration is embargoed and holds only the hashes of the three frozen plan documents, so no third-party timestamp precedes the run start. The repeat of 8.11 on held-out units and the per-fold recomputation of C01_E02 used held-out data after the single confirmatory run; the specification says the held-out split runs once, so they are departures from that wording, and neither can change an outcome. Known limitation: three inputs that fix the unit set and the age subset (DIV table, session index, unit list) were not hashed by the guard; their hashes were recorded on 29 Oct and the two tracked ones are unchanged since the tag. On 2026-11-02 the SHA-256 of every frozen file, the split file, the environment lock and the gate results were recomputed by C.C. on a fresh unzip of the code snapshot and match the hashes written by the run; W.R. confirmed the hashes on a separate fresh unzip the same day. On 2026-11-04 the public code repository (https://github.com/amisepa/cortical-anticipation-001611) was created from the same files; its check script (check_hashes.py) reports every frozen file identical to the run record.

